# circASbase: A Comprehensive Database of Alternative Splicing Events in circRNAs

**DOI:** 10.1101/2025.03.09.642279

**Authors:** Lingxiao Zou, Jian Zhao, Haojie Li, Chen Xu, Yulan Wang, Xuejiang Guo, Xiaofeng Song

**Affiliations:** Department of Biomedical Engineering, Nanjing University of Aeronautics and Astronautics, Nanjing 211106, China; State Key Laboratory of Reproductive Medicine and Offspring Health, Department of Histology and Embryology, Nanjing Medical University, Nanjing 211166, China; Changzhou Medical Center, The Affiliated Changzhou Second People’s Hospital of Nanjing Medical University, Changzhou Second People’s Hospital, Nanjing Medical University, Changzhou 213000, China; Medical Research Center, Changzhou Maternal and Child Health Care Hospital, Changzhou Medical Center, Nanjing Medical University, Changzhou 213000, China

**Keywords:** Circular RNA, Alternative splicing, Full-length isoform, Coding potential, Database

## Abstract

Despite extensive studies highlight the critical roles of alternative splicing in generating mature circRNA isoforms and enhancing their function diversity, a significant gap remains in the availability of dedicated databases for circRNA alternative splicing events. To bridge this gap, we developed circASbase, a pioneering and comprehensive database that catalogues 452,129 alternative splicing events in 884,047 full-length circRNAs from 581 samples across 13 species, and provides rich annotations to facilitate understanding the splicing regulation of circRNAs. Our findings reveal substantial differences between circRNAs and linear transcripts regarding the distribution and occurrence of alternative splicing events, highlighting the unique regulatory landscape of circRNAs. These unique splicing events result in functional differences of circRNAs by affecting IRES sites, m6A sites, ORFs, protein features, miRNA targets, and more. In summary, circASbase not only covers the urgent need of the research community for data repositories, but also represents a significant advancement in our understanding of circRNA biology. With its user-friendly interfaces and web-based visualization tools, circASbase is poised to become an indispensable resource for researchers exploring the regulatory mechanisms and functional roles of alternative splicing events in circRNAs. This database will continuously drive new insights and discoveries in the field, setting the stage for further advancements in circRNA research. circASbase is available at http://reprod.njmu.edu.cn/cgi-bin/circASbase/

## Background

Circular RNAs (circRNAs) are a novel class of functional RNAs characterized by covalently closed-loop structures generated through back-splicing [1]. They are ubiquitously present in eukaryotes and exhibit tissue- and developmental stage-specific expression patterns [2–4]. Initially, circRNAs were thought to result from erroneous splicing, but Memczak et al. revealed that circRNAs act as sponges for microRNAs (miRNAs) [5–7]. Currently, circRNAs are recognized for their roles in diverse biological processes, including cell proliferation, migration, metastasis, and apoptosis [8]. Furthermore, growing evidence has revealed that dysfunction of circRNAs is closely associated with various diseases, particularly cancers [9, 10].

Most circRNAs are originated from pre-mRNAs and typically consist of multiple exons. While circRNAs were initially distinguished from their linear counterparts by the back-splice junction (BSJ) sites, it is now known that multiple circRNAs can share the same BSJ via alternative splicing [11]. Furthermore, recent studies indicate that circRNAs can be spliced distinctly from their linear counterparts [12, 13]. Notably, several exons or introns have been found to be preferentially included in circRNAs through alternative splicing mechanisms [14]. Despite the discovery and investigation of a large number of circRNAs based on the recognition of back-spliced junctions, our understanding of alternative splicing events in circRNAs remains limited.

Alternative splicing of mRNA and lncRNA has been extensively studied, and the association of RNA splicing regulation with various biological processes, such as sex determination, T-cell activation, and synapse development, has been well-established [15–17]. Aberrant splicing has been associated with aging and various diseases, including cancers, inflammation, and metabolic disorders [18]. Similar to mRNA and long noncoding RNA (lncRNA), the function of circRNA also relies primarily on its sequence, which determines miRNA-sponge activity, interaction with protein, and coding potential, etc [19]. Dysregulation of alternative splicing can change circRNAs’ roles in biological processes, leading to dysfunction and disease [20, 21].

Advances in RNA sequencing and analysis have led to the discovery of an increasing number of circRNAs. To deposit and annotate these circRNAs, multiple general and specialized databases have been developed, such as circBase, CircNet, CIRCpedia, circRNADb, CSCD, Circbank, and circAtlas [2, 22–27]. Most of these databases use the BSJ as the unique identifier for circRNAs and generate their sequences by sequentially assembling exons located between the 5’ and 3’ ends of the BSJ. However, only a few databases, such as circAtlas, PlantcircBase, and isoCirc, provide full-length circular RNA isoforms, which are essential for inferring alternative splicing events within circRNAs [26, 28, 29].

Recently, several methods have been proposed to assemble full-length circRNA isoforms. Zheng et al. introduced CIRI-full that utilizes the back-splice junction and reverse overlap to reconstruct circular isoforms, which is based on a particular library preparation strategy [30]. Wu et al. developed CircAST to assemble alternatively spliced circRNA transcripts with RNase R-treated RNA-seq data [31]. Xin et al. reported isoCirc for sequencing full-length circRNA isoforms using rolling amplification and long-read sequencing strategy [28]. While thousands of alternative splicing events within the internal part of circRNAs have been identified, there is currently no comprehensive database for depositing and annotating these splicing events. Here, we developed circASbase, a comprehensive database of alternative splicing events in circRNAs. circASbase provides comprehensive annotations for each splicing event, detailing its effect on circRNAs’ coding potential and regulatory features, such as internal ribosome entry sites (IRES), N6-methyladenosine (m6A) sites, open reading frames (ORFs), protein features, miRNA targets, and more.

## Construction and content

### Construction and functionalities of the circASbase

To develop a high-quality circRNA database, we queried the SRA database (https://www.ncbi.nlm.nih.gov/sra/) using two keywords until January 21, 2022. Our database includes only RNA-seq data processed by RNase R, an exoribonuclease enzyme capable of efficiently degrading linear RNA to enrich circular RNA [32, 33]. A total of 581 samples were collected (**Table S1**), including 13 species (human, mouse, rat, macaque, rabbit, cattle, pig, chicken, zebrafish, honey bee, fruit fly, thale cress and rice). The structure and content of circASbase are depicted in **Figure 1**. Briefly, all circRNAs were identified using CIRI2 [34], except for circRNAs collected from other open-source projects. Full-length sequences were assembled with CircAST [31]. For circRNAs not processed by CircAST, full-length sequences were assembled based on reference genomes and genome annotation files (**Table S1**). These full-length sequences are also annotated for components and internal structures. Potential alternative splicing events were detected using chromosomal coordinates and internal structures. In addition, functional elements such as internal IRESs, ORFs, m6As, miRNA targets, protein glycosylation sites and phosphorylation sites on full-length circRNAs or their coding proteins were predicted. Sequence conservation was evaluated using PhastCons and PhyloP models [35, 36]. Finally, we developed circASbase with dynamic visualization and database management features, offering search, browse, and download capabilities.

**Figure 1.**
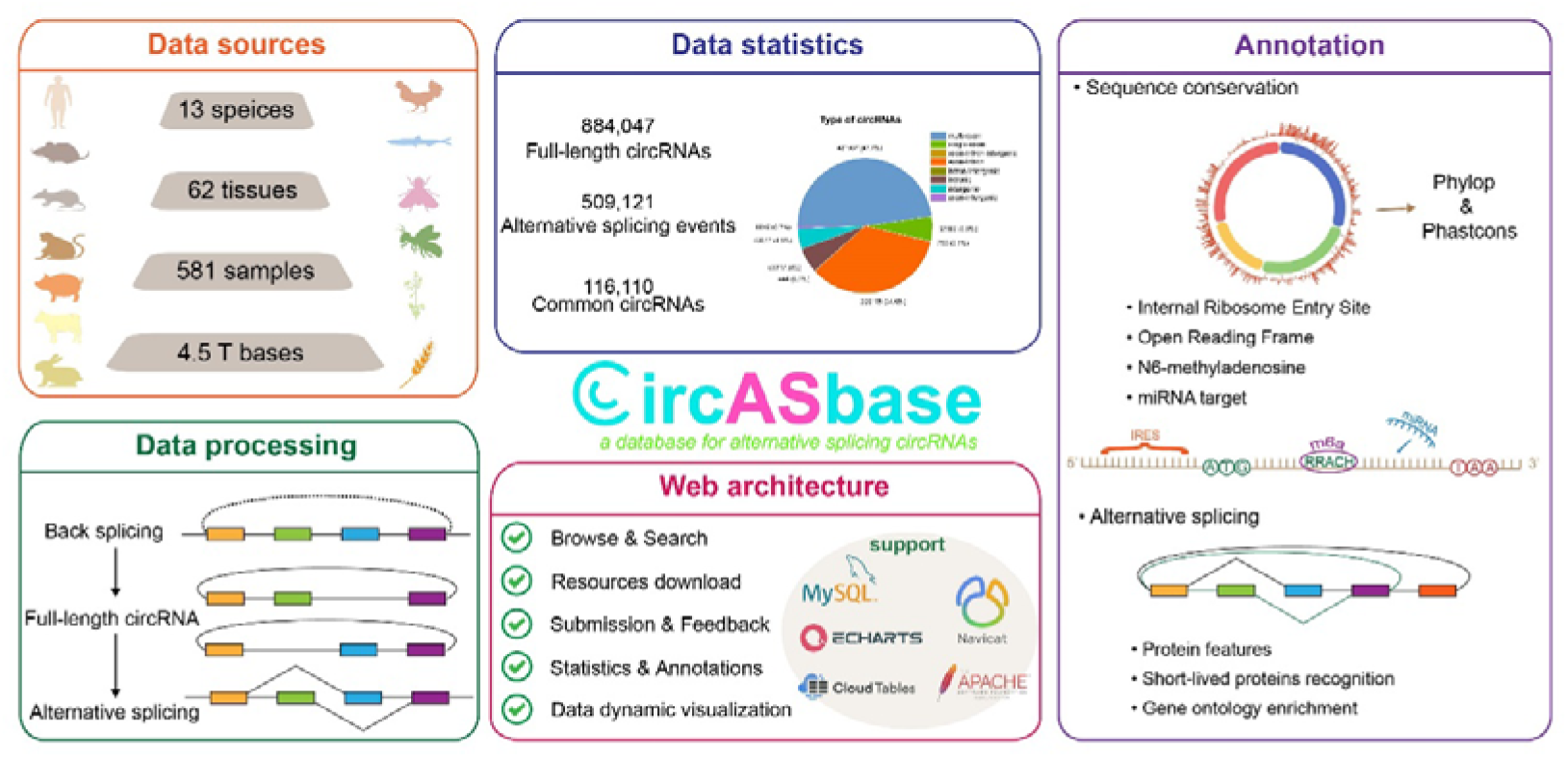
Database construction overview of circASbase. Process of data collection, full-length circRNAs prediction, alternative splicing events identification, circRNAs annotation and webpage construction.

### Full-length assembly of circRNAs

We employed the BWA-MEM algorithm to align reads to the reference genome [30]. Subsequently, CIRI2 was used to identify back-splice junctions (BSJ), and CircAST reconstructed the internal structures of circRNAs [31, 34]. This pipeline enabled us to obtain numerous high-confidence full-length circRNA sequences (**Supplementary material: Methods**). Since CircAST is an exon annotation-based approach to constructing splicing graphs, it may lose some circRNAs composed of intronic sources or intergenic sequences. For BSJ sites that cannot be assembled by CircAST, we directly concatenated exons between the back-splice regions (**Figure S1A**). This may result in the inclusion of introns and intergenic regions in circRNAs, as BSJ sites were not always located in the exonic regions.

Additionally, we collected full-length circRNA sequences from open-source projects or databases, comprising 286,696 for humans (**Figure S1B**), 6519 for rice (including 466 validated), and 4663 for thale cress (including 137 validated) [29, 37]. To eliminate redundancy, we used the CD-HIT-EST algorithm to remove circRNAs with over 95% sequence identity [38].

### Alternative splicing events in circRNAs

We annotated circRNA isoforms for each species in GTF format and devised a strategy to identify and classify alternative splicing events in circRNAs by integrating our annotations with genome annotation files from the Ensembl database. Our strategy categorizes alternative splicing into seven types: A3 (alternative 3’ splice site), A5 (alternative 5’ splice site), ES (exon skip), EES (exclusive exon skip), IR (intron retain), ABS3 (alternative 3’ back-splice site), and ABS5 (alternative 5’ back-splice site).

### Sequence conservation analysis

PhyloP evaluates only the current column of the alignment, while PhastCons also takes neighboring columns into account, making PhastCons more sensitive to detecting conserved regions and PhyloP more effective at defining them [35]. Using the chromosomal position of circRNAs, we extracted sequence conservation scores (PhastCons and PhyloP) from UCSC database with bwtool [36]. Since conservation data were available only for human, mouse, rat, and chicken, our conservation analysis focused on these species. CircRNAs with an average PhastCons score >0.6 and an average Phylop score >2 [39, 40] were considered highly conserved.

### Large-scale functional annotation of the circRNAs

We used IRESfinder and SRAMP tools to identify potential IRES and m6A sites in circRNAs, respectively [41, 42]. To determine the longest ORF in circRNAs, we first replicated and concatenated the circRNA sequence three times to simulate circular reading frame. Then, we extracted all start and stop codons from the amplified sequence and selected the farthest pair as the ORF, which allows for overlapping ORFs in circRNAs that undergo looping. Approximately 60% ORFs of circRNAs cross the BSJ site (**Figure S3**). Using SProtP [43], we predicted the half-lives of proteins encoded by ORFs in human circRNAs. For glycosylation site prediction, we used NetCGlyc for C-mannosylation, NetNGlyc for N-linked glycosylation, and NetOGlyc for O-GalNAc glycosylation [44–46]. We also employed the GPS (v2.1) tool, using a high-threshold model, to predict phosphorylation sites [47]. To identify potential miRNA binding sites in circRNAs, we used TargetScan, miRanda, and RNAhybrid to identify potential miRNA binding sites in circRNAs [48–50], with mature miRNA sequences sourced from the miRBase database.

### Database construction and implementation

The core framework of circASbase utilizes Apache as the web server and MySQL as the database server. All data are managed within the MySQL database system. To enhance user experience, we developed the interface primarily in JavaScript, and implemented tables using DataTables jQuery plugin (version 1.11.4). For data visualization, we employed ECharts (version 5.2.1), a JavaScript library. The web services of circASbase have been tested on several popular browsers, including Google Chrome, Edge, and Firefox.

## Utility and discussion

Here, we innovatively developed an interactive database that integrates alternative splicing and coding potential data for full-length circRNAs. This is the first comprehensive database to include full-length circRNAs from animals, insects, and plants. To construct this database, we collected 581 samples from 13 species, processed approximately 4.5 terabytes of data, and incorporated circRNA sequences from external sources. In total, we identified 884,047 full-length circRNAs, with 452,129 alternative splicing events (**Figure 2C**, **3A**) detected. Additionally, through homology and sequence similarity analysis, we identified 116,110 conserved circRNAs (**Figure 4A**) shared by at least two species.

**Figure 2.**
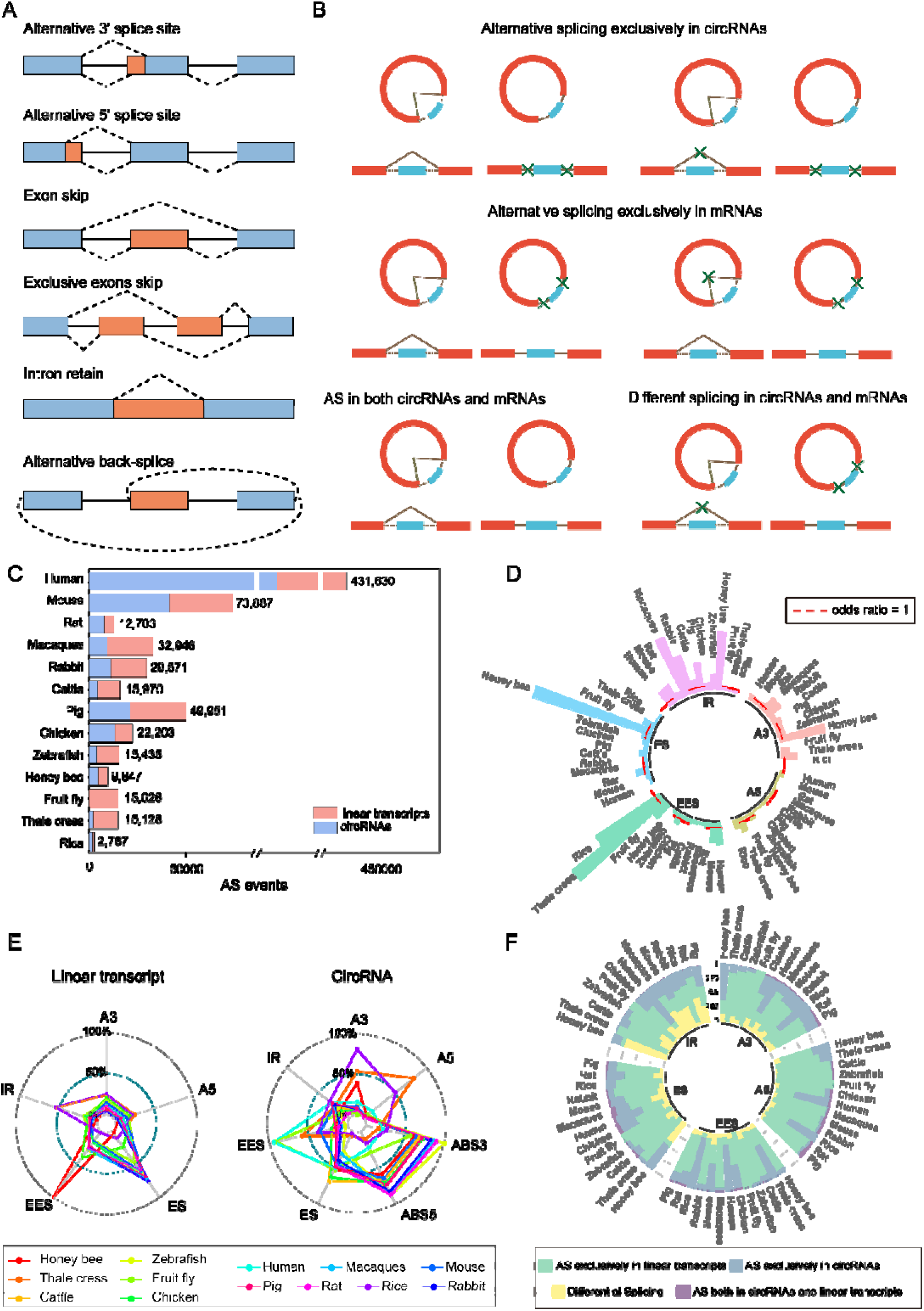
Statistics and analysis of alternative splicing events. (A) Classification diagram of alternative splicing events. (B) Classification diagram of different alternative splicing events between circRNAs and mRNAs. The classification theme is that when two splicing patterns occur in the same region of circRNAs, they can be counted as alternative splicing events of circRNAs. The same with mRNAs. (C) Statistics of alternative splicing events in circRNAs and mRNAs of various species. (D) Distribution of different types of alternative splicing events in linear transcripts and circRNAs from 13 species. (E) Odds ratios of various types of alternative splicing in linear and circular RNAs. The odds ratio reflects the preference of a certain type of alternative splicing events in circRNAs and mRNAs. (F) Statistics of different alternative splicing events in circRNAs and mRNAs. CircRNA-specific or mRNA-specific alternative splicing events account for the majority, and shared alternative splicing events are still rare.

### Alternative splicing events in circRNAs

In this study, we comprehensively examined alternative splicing events in covalently closed circRNAs, which lack free 3’ or 5’ ends. A total of 452,129 alternative splicing events were identified across circRNAs from 13 species (**Figure 2C**). Of these, 329,269 events were observed in human, whereas only 43 events were detected in fruit fly. It is important to note that the differences in number of events may reflect variations in sample sizes rather than species-specific phenomena. For a detailed summary of these results, refer to **Table S2**.

We further analyzed the distribution of alternative splicing events in circRNAs and found that EES was the most common event, accounting for 30.1% of cases. This was followed by ES (20.1%), ABS5 (14.4%), ABS3 (14.3%), and IR (11.3%) (**Table S2**). Based on these splicing events, we obtained 2,372,838 circRNA pairs (presence of one or more AS events between two circRNAs) and assessed the effect of different splicing patterns on the functional elements of circRNAs. The results indicated that for most of circRNA pairs, different splicing patterns resulted in alterations in ORFs, IRESs, and m6a sites (**Figure S1D-S1F**), which may affect the function of circRNAs [51, 52].

### Comparison of alternative splicing between circRNAs and their corresponding linear transcripts

The identification of alternative splicing events in circRNAs enables a comparative analysis of splicing event distributions between circRNAs and their corresponding linear transcripts across multiple species (**Table S3**). Our study identified 452,129 alternative splicing events in circRNAs and 275,001 in their linear counterparts across 13 species (**Figure 2A**, **2C**). Interestingly, in linear transcripts, ES is the predominant splicing event in most species, except for insects (e.g., honey bee and fruit fly), where EES is most prevalent, and in plants, where IR is the dominant event **(Figure 2E)**. The proportions of A3 and A5 events are similar across all species.

In contrast, circRNAs exhibit distinct patterns of alternative splicing. ABS3 and ABS5 events are specific to circRNAs and are abundant across most species. To further explore the differences in splicing events between circRNAs and linear transcripts, we calculated the odds ratios for the five types of splicing events common to both RNA types (**Figure 2D**). Our analysis revealed a higher incidence of IR events in circRNAs, with most species displaying a similar distribution of splicing events between circRNAs and linear transcripts. However, circRNAs exhibit a higher frequency of splicing events in plants and insects compared to their linear transcripts. In summary, our findings highlight significant interspecies variation in the distribution of splicing events between circRNAs and linear transcripts.

To further characterize the differences in alternative splicing between circRNAs and linear transcripts, we classified the events into four types based on full-length sequences: (1) AS events present in both circRNAs and linear transcripts, (2) AS events exclusively in circRNAs, (3) AS events exclusively in linear transcripts, and (4) different splicing patterns between circRNAs and linear transcripts. (**Figure 2B**). The key criterion in this classification is whether the two splicing modes corresponding to an alternative splicing event occurs in both circRNAs and their corresponding linear transcripts. Although the proportions of these types vary across the five event categories (A3, A5, ES, EES, IR), 98% (616,816/629,298) events are not shared between circRNAs and linear transcripts (**Figure 2F**, **S2**), with shared events being relatively rare. In addition, we assessed the conservation between circRNA-exclusive and mRNA-exclusive fragments in splicing events. We found that circRNA-exclusive splicing fragments were more conserved than mRNA-exclusive splicing fragments (**Figure S3A, S3B**).

### Structural characteristics and conservation of circRNAs

Previous research has demonstrated that circRNAs vary widely in length, with exonic circRNAs ranging from 200 bp to 1,000 bp, and intergenic circRNAs typically exceeding 3,000 bp [53, 54]. We analyzed the length distribution of circRNAs across 13 species (**Figure 3B**) and found that in most animals, circRNAs are less than 3,000 bp. The length distribution in other animal species resembles that of humans, though more dispersed. Notably, plant circRNAs are significantly shorter than those of animals, predominantly between 200 and 600 bp. In contrast, insect circRNAs are longer and more dispersed compared to other species (**Figure 3B**). These findings suggest substantial interspecies variation in circRNA lengths, particularly among animals, plants, and insects.

**Figure 3.**
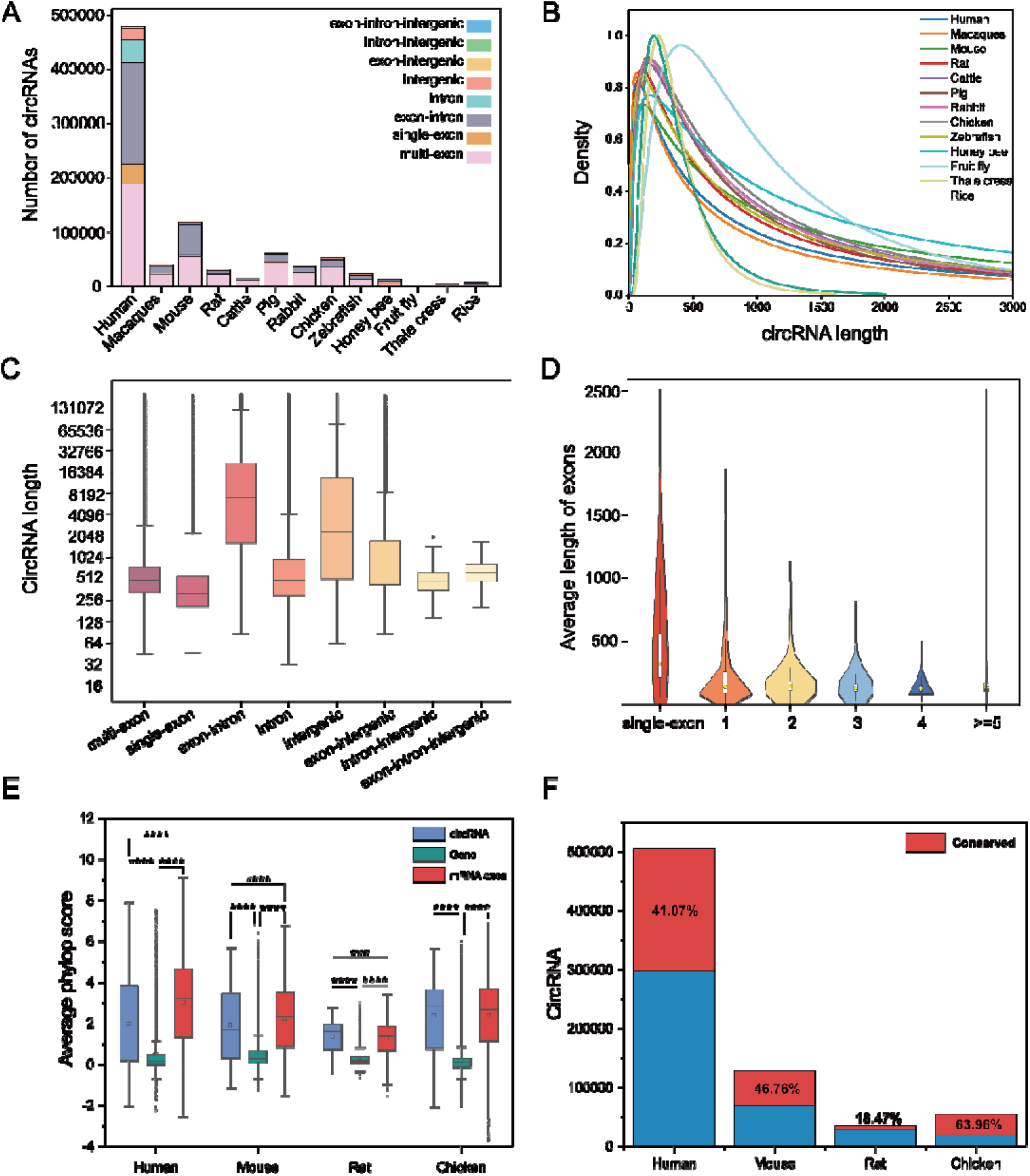
Structural characteristics and conservation of circRNAs. (A) Number of circRNAs per species and distribution of different types. (B) Densitometric distribution of circRNAs length in each species. (To exclude the effect of ultra-long circRNAs on the graphs, only data in the range of 0-3,000 were intercepted). (C) Boxplot for length distribution of different types of circRNAs. (D) Average length of exons in different types of circRNAs. (E) Average phylop scores of circRNAs, genes and mRNA exons (Student’s t-test, *:P<0.05, **:P<0.01, ***:P<0.001, ****:P<0.0001). (F) Proportion of highly conserved circRNAs (average PhastCons score >0.6, average Phylop score >2).

Given that length differences may be attributed to the internal structure of circRNAs, we categorized them into eight types: single-exon, multi-exon, exon-intron, intronic, intergenic, exon-intergenic, intron-intergenic, and exon-intron-intergenic. Approximately half of these circRNAs consists of multiple exons (multi-exon), and more than one-third of circRNAs contains exons and introns (exon-intron). Roughly 6% circRNAs consist of a single exon (single-exon), and about 89% circRNAs contain 1 to 5 exons. Conversely, around 11% of circRNAs are entirely intronic or intergenic, with this subset primarily observed in human (**Figure 3A**). Furthermore, several circRNAs (exon-intergenic, intron-intergenic, exon-intron-intergenic) cross the known boundaries of their host genes.

Most circRNAs fall within the 500 to 800 bp range (**Figure 3C**), with intron retention significantly increasing their length. Notably, the length difference between single-exon circRNAs and other exonic circRNAs is smaller than expected (**Figure 3C**), suggesting that single-exon circRNAs may have longer exons. To better understand their differences, we calculated the average exon length and number for each circRNA type. The result revealed that exons in single-exon circRNAs are substantially longer than those in other circRNA types (**Figure 3D**), corroborating previous findings [37, 55, 56].

Moreover, we examined the conservation of circRNAs. Based on PhyloP and PhastCons scores, circRNAs are slightly less conserved than exons of mRNAs, but more conserved than entire genes (**Figure 3E**, **S3C**). We considered circRNAs with average Phylop score >2 and average PhastCons score >0.6 as highly conserved, with approximately 40% of circRNAs highly conserved (**Figure 3F**). Additionally, the regions near back-splice junction (BSJ) sites exhibited remarkable conservation (**Figure S3D-S3G**).

In addition, we compared circRNAs from four shared species in circASbase and circAtlas 3.0, and 56.7% of BSJs (human: 47.2%, mouse: 70.4%, rat: 70.5%, pig: 87.0%) from circASbase were the same as those in circAtlas (**Figure S4A**). For the circRNAs with the same BSJs, we compared their full-length sequences in the two databases and found 87,107 circRNAs (human: 43,414, mouse: 19,554, rat: 7,873, pig: 16,266) that were completely identical (Similarity is 100%) in both (**Figure S4B**).

### Identification of common circRNA isoforms derived from orthologous genes

Recent studies suggest that circRNAs are closely associated with species identity [6, 57–59]. Orthologous genes producing circRNAs generally possess more conserved splicing sites and exons than other genes [60]. To explore conserved circRNA isoforms and understand their genetic basis through evolutionary analysis, we conducted a sequence similarity analysis of circRNA isoforms generated by orthologous genes. A total of 116,110 eligible circRNAs were found in at least two of nine species (**Figure S4C**). In addition, 646 circRNAs were expressed across all seven mammalian species (**Figure 4A**). These findings indicate that the presence of shared circRNAs between species is a common phenomenon. Furthermore, species that are evolutionarily closer, such as human and macaque, or mouse and rat, tend to share a higher number of circRNAs.

**Figure 4.**
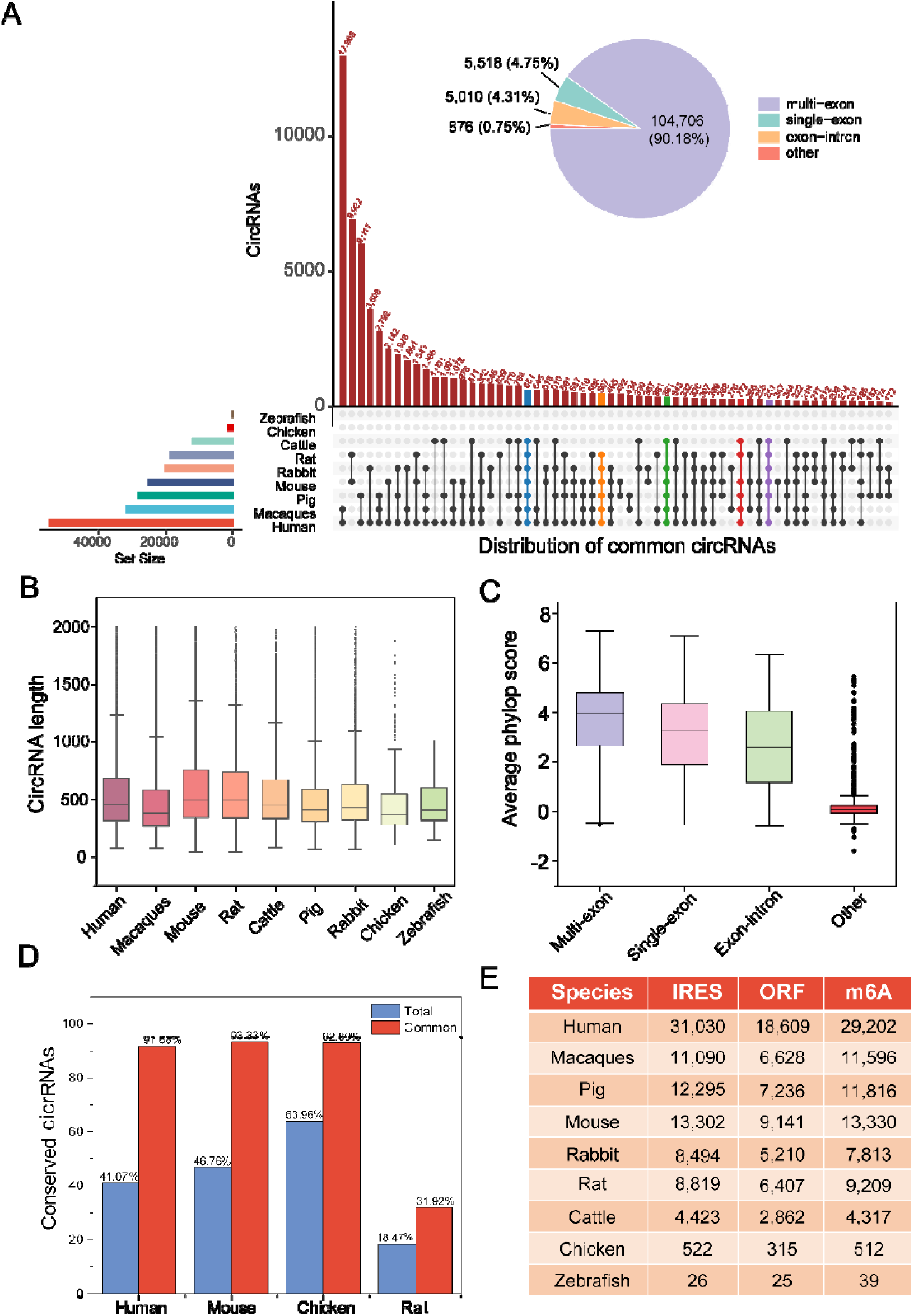
Characterization of common circRNAs in nine animals. (A) An UpSetR plot of the intersection between sets of common circRNAs derived from at least two species. Horizontal bar graphs indicate the number of common circRNAs in each species, and the connecting lines indicate different patterns shared among species; patterns shared in more than five species have been labeled in color. Vertical bar graphs indicate the number of common circRNA pairs in the corresponding sharing mode. (B) Length distribution of common circRNAs in each species. (C) Conservation assessment (phylop) of common circRNAs of different types. (D) Proportion of highly conserved circRNAs in two groups. (E) Statistics of common circRNAs containing potential coding elements in each species.

Similarly, we investigated structural characteristics of circRNAs common in different species and found that the proportion of multi-exon circRNAs was significantly increased than the proportion of multi-exon circRNAs among all circRNAs (**Figure 4A**). The circRNAs common in different species contain very few introns and intergenic regions, which leads to further compression of the length range (**Figure 4B**). The circRNAs common in different species were in a highly conserved state (average Phylop score >2 and average PhastCons score >0.6), although the scores varied slightly (**Figure 4D**, **S4D-S4E**). The existence of introns and intergenic regions can significantly reduce the conservation level of circRNAs common in different species (**Figure 4C**, **S4F**). Since circRNAs existing commonly in two or more species are mostly conserved, they are likely to play important functional roles. Almost all these common circRNAs are annotated with ORFs, IRESs and m6A sites (**Figure 4E**), which means they may have strong coding potential.

### Browse and search

CircASbase is a user-friendly database designed to support searching, browsing, and downloading of circRNA data. It utilizes web visualization technology to display alternative splicing events in circRNAs. Users can access the database in two ways: (1) by selecting a target species in the search box on the homepage and searching with specific keywords (e.g., circRNA ID, gene ID, etc.) (**Figure 5A**); or (2) by using the advanced search page, where queries can be refined with keywords such as species, circRNA ID, gene ID, chromosome, strand, circRNA type, and sample name (**Figure 5B**). The query results are displayed in paginated tables, allowing users to further sort and filter the data by specified conditions (**Figure 5C**).

**Figure 5.**
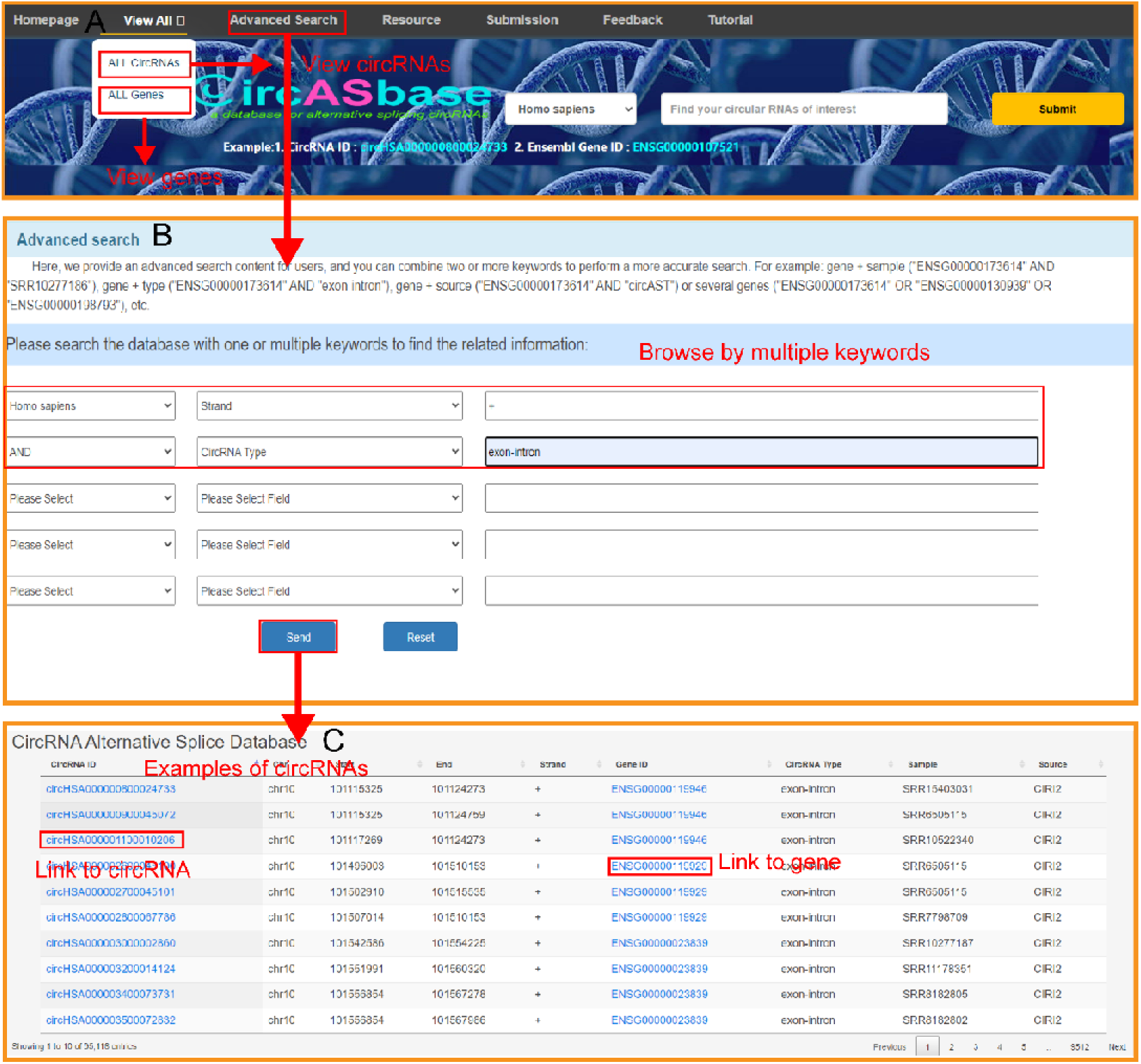
Screenshot of database browse. (A) Navigation bar and home page search section of the database, circRNAs and genes could be separately browsed in the ‘View All’ section or retrieved in the search box. (B) Advanced search function. Search by combination of multiple keywords. (C) Options for customized browsing of retrieved circRNAs and genes.

### Data visualization

To improve the usability of circASbase, alternative splicing events in circRNAs from 12 different species (excluding honey bee) are presented graphically. The gene page displays the genomic structure in the top panel (**Figure 6A**), followed by diagrams of circRNA isoforms derived from the gene, with exons color-coded to match the genomic structure (**Figure 6B**). A search box in the upper left corner allows users to select specific circRNA isoforms by their IDs or back-splicing sites (**Figure 6C-6D**). In addition to the genomic structure of circRNA isoforms, diagrams illustrating alternative splicing events between isoforms are provided. Users can select specific splicing types to view, gaining a more intuitive understanding of the AS events (**Figure 6E**). Additionally, the database offers detailed pages for each circRNA isoform, displaying graphical information on sequence conservation, ORFs, IRESs, and m6A sites (**Figure 7**).

**Figure 6.**
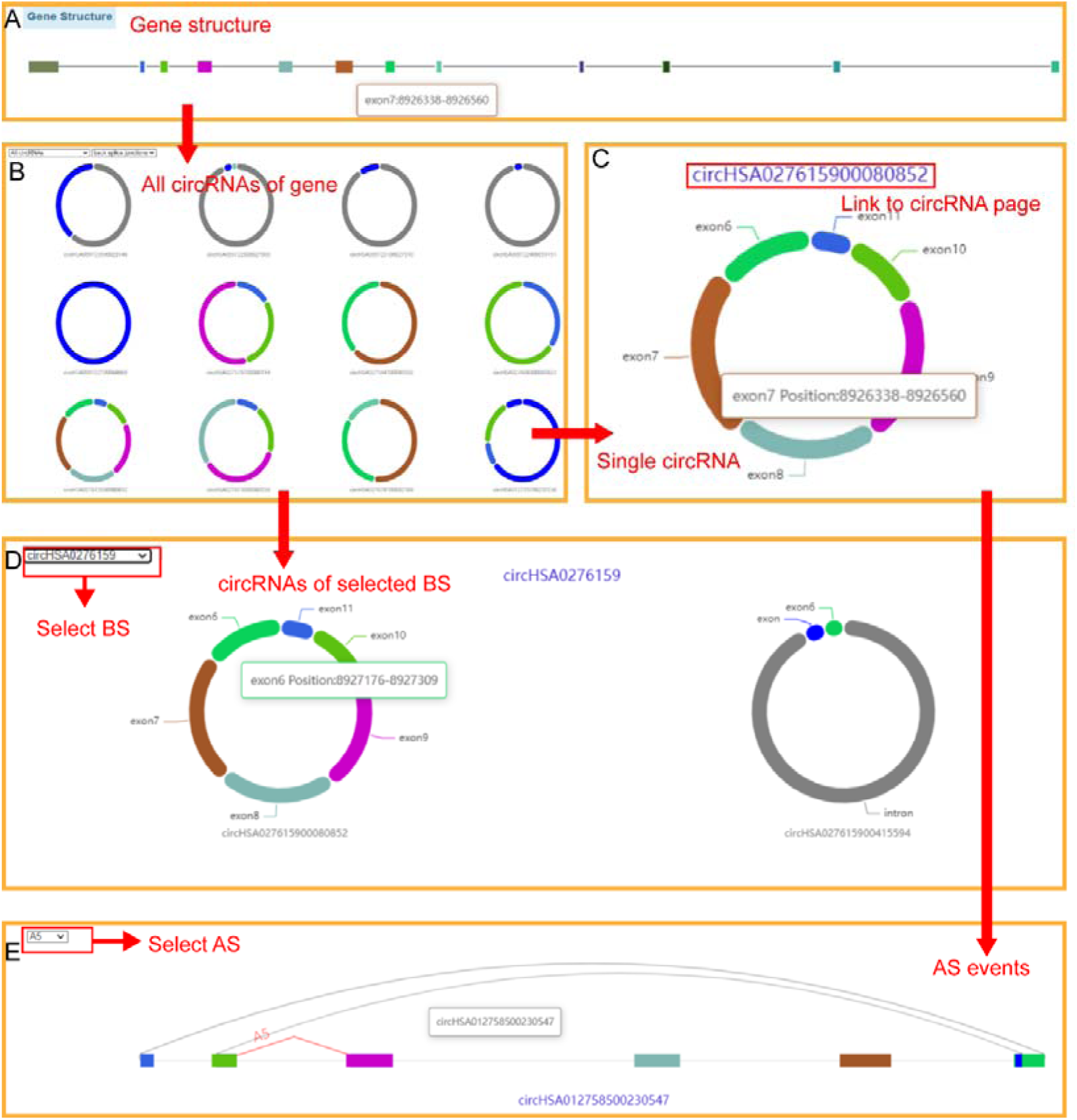
Page structure of alternative splicing events. (A) Display of host gene structure. (B) Charts of all circular isoforms of the queried gene. (C) Schematic diagram of a single circRNA. The link can jump to the corresponding circRNA page. (D) CircRNAs with same back-splice junction site. (E) Visualization of alternative splicing events associated with selected circRNA.

**Figure 7.**
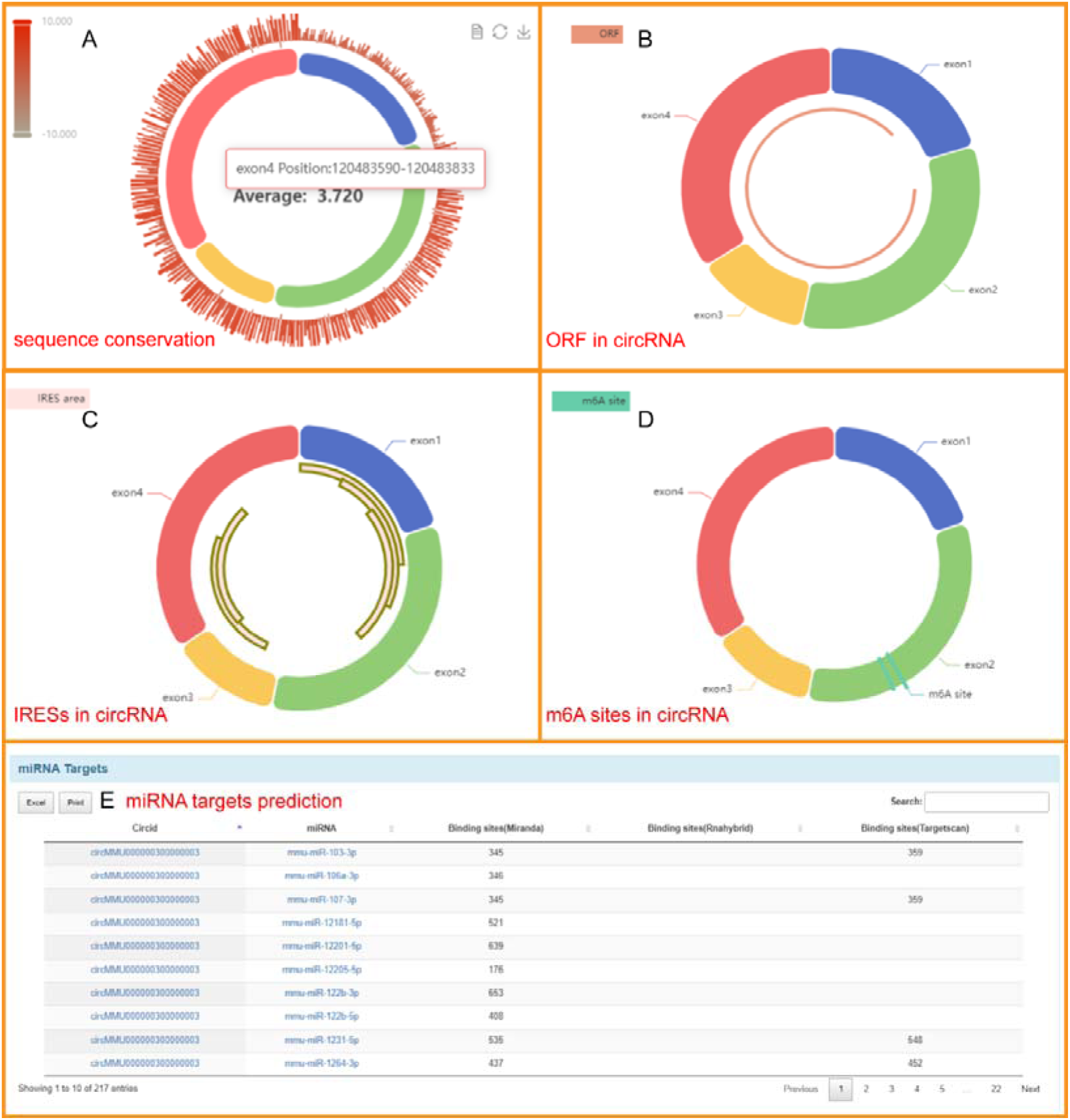
Overview of circRNA information page. (A) Schematic diagram of sequence conservation for the queried circRNA. (B) Schematic diagram of the open reading frame in the queried circRNA. (C) Schematic diagram of internal ribosome entry sites in the queried circRNA. (D) Schematic diagram of N6-methyladenosine sites in the queried circRNA. (E) Distribution table of microRNA targets in the queried circRNA.

## Conclusions

In this study, we assembled 884,047 full-length circRNA isoforms with splicing annotations from 13 species. Our analysis revealed that the length distribution of circRNAs differs significantly at the kingdom level, with plant circRNAs being notably shorter than those of animals. Moreover, the exons of single-exon circRNAs are significantly longer than those of other circRNA types. Additionally, we identified 116,110 orthologous circRNAs common in at least two species. We also observed that the alternative splicing events of circRNAs and linear transcripts differ markedly in both splicing forms and preferences, with few shared alternative splicing events.

In circASbase, we provide dynamic visualizations of all alternative splicing events. The database also includes detailed information on circRNAs, such as general information, full-length sequences, IRESs, m6A sites, ORFs, phosphorylation sites, protein glycosylation sites, conservation scores and miRNA targets. Compared to existing circRNA databases, circASbase offers large-scale visualization of alternative splicing events across 13 species from three kingdoms (animals, insects, and plants). In conclusion, circASbase serves as a valuable resource for understanding the mechanisms of alternative splicing in circRNAs and their functional roles.

## Availability

CircASbase is freely accessible at http://reprod.njmu.edu.cn/cgi-bin/circASbase/.

## Authors’ contributions

XJG and XFS conceived and designed the study, revised the manuscript. LXZ, JZ and HJL carried out data acquisition and analysis. LXZ performed website construction and drafted the manuscript. YLW and BJ revised figures, provided scientific advice and contributed to result interpretations. XJG and XFS assisted with manuscript review and revision. All authors read and approved the final manuscript.

## Acknowledgements

The study was supported by the Chinese National Natural Science Foundation (Grants No. 62273175 to XF, No. 82221005, 82371606 to XG, No. 62003165 to JZ), the National Key R&D Program of China (2021YFC2700200 to XG), the Key R&D projects of Jiangsu Province (No. BE2022843 to XF), the Scientific Research Project of Changzhou Medical Center of Nanjing Medical University (CMCP202303 to HL), the Jiangsu Funding Program for Excellent Postdoctoral Talent (2023ZB319 to HL), the China Postdoctoral Science Foundation (2024M750276 to HL).

