## supplementary material for "circASbase: A Comprehensive Database of Alternative Splicing Events in circRNAs"

| Species | # Full-length sequences | # Samples | # Reference genome |
| --- | --- | --- | --- |
| Homo sapiens  Mus musculus  Rattus norvegicus  Macaca mulatta  Oryctolagus cuniculus  Bos taurus  Sus scrofa  Gallus gallus  Danio rerio  Apis mellifera  Drosophila melanogaster  Arabidopsis thaliana  Oryza sativa | 478460  118728  31085  40131  38232  14830  61432  52593  22669  12340  296  5503  7748 | 299  81  9  61  9  8  6  50  17  39  2  Collected  Collected | hg19  mm10  rn6  rheMac10  oryCun2  bosTau9  susScr11  galGal6  danRer11  apiMel2  dm6  TAIR10  IRGSP-1.0 |

**Table S1.** The statistics of full-length circRNA sequences and their corresponding samples.

| Species | #A3 | #A5 | #ABS3 | #ABS5 | #ES | #EES | #IR | Total |
| --- | --- | --- | --- | --- | --- | --- | --- | --- |
| H. sapiens | 20360 | 17301 | 29079 | 29763 | 67798 | 124940 | 40028 | 329269 |
| M. musculus | 1582 | 370 | 10819 | 11049 | 7756 | 5977 | 4149 | 41702 |
| R. norvegicus | 150 | 26 | 2902 | 2819 | 1523 | 380 | 200 | 8000 |
| M. mulatta | 208 | 35 | 3405 | 3407 | 1241 | 323 | 1100 | 9719 |
| O. cuniculus | 260 | 43 | 3807 | 3807 | 1874 | 449 | 1087 | 11327 |
| B. taurus | 79 | 21 | 1194 | 1158 | 1213 | 349 | 316 | 4330 |
| S. scrofa | 557 | 177 | 6168 | 6367 | 4284 | 1399 | 2365 | 21317 |
| G. gallus | 275 | 66 | 3947 | 3883 | 3464 | 1150 | 715 | 13500 |
| D. rerio | 158 | 26 | 1603 | 1395 | 455 | 62 | 287 | 3986 |
| A. mellifera | 781 | 78 | 1241 | 1188 | 1035 | 310 | 307 | 4940 |
| D. melanogaster | 0 | 1 | 10 | 5 | 9 | 16 | 2 | 43 |
| A. thaliana | 509 | 755 | 154 | 107 | 56 | 562 | 223 | 2366 |
| O. sativa | 530 | 328 | 176 | 106 | 25 | 253 | 212 | 1630 |

**Table S2.** Distribution of different types of circRNA splicing events in 13 species.

| Species | #A3 | #A5 | #ES | #EES | #IR | Total |
| --- | --- | --- | --- | --- | --- | --- |
| H. sapiens | 9609 | 9398 | 55290 | 21649 | 6415 | 102361 |
| M. musculus | 3989 | 4014 | 16985 | 4331 | 2866 | 32185 |
| R. norvegicus | 401 | 362 | 2438 | 1080 | 422 | 4703 |
| M. mulatta | 732 | 960 | 14621 | 5491 | 1423 | 23227 |
| O. cuniculus | 559 | 561 | 11998 | 3926 | 1200 | 18244 |
| B. taurus | 572 | 648 | 6433 | 2838 | 1149 | 11640 |
| S. scrofa | 1376 | 1742 | 17240 | 5941 | 2335 | 28634 |
| G. gallus | 381 | 434 | 5468 | 1260 | 1163 | 8706 |
| D. rerio | 1707 | 1730 | 4734 | 1490 | 1788 | 11449 |
| A. mellifera | 298 | 150 | 165 | 4216 | 78 | 4907 |
| D. melanogaster | 2627 | 2474 | 3151 | 5118 | 1656 | 15026 |
| A. thaliana | 2667 | 2736 | 1182 | 264 | 5913 | 12762 |
| O. sativa | 249 | 227 | 114 | 23 | 544 | 1157 |

**Table S3.** Distribution of different types of linear transcripts splicing events in 13 species.

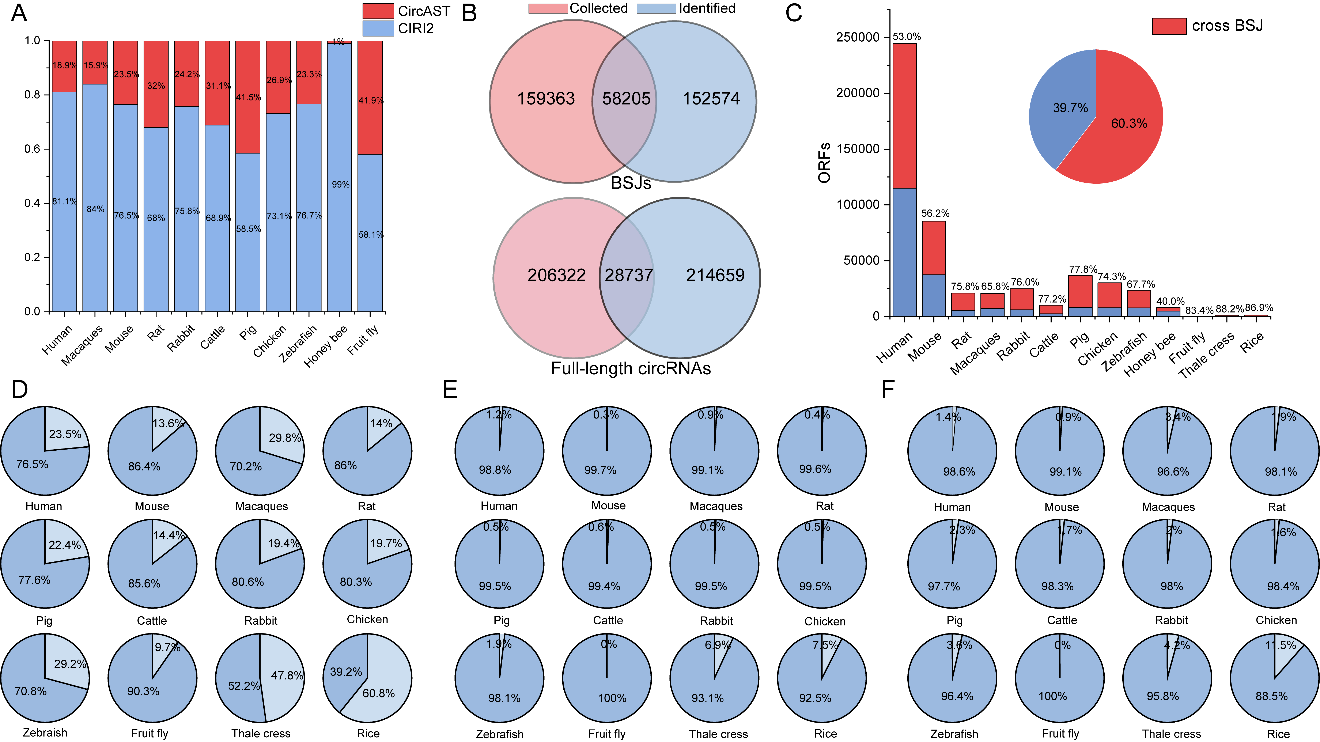

**Figure S1.** (A) Proportion of full-length circRNAs assembled by CircAST or CIRI2 after removal of redundancy. (B) Overlapped BSJs and full-length circRNAs of human. (C) Statistics on ORFs of circRNAs crossing the BSJ sites in 13 species. (D) Effect of AS events on ORFs in each circRNA pair. Darker color indicates ORF changed, lighter color indicates ORF not changed. (E) Effect of AS events on IRESs in each circRNA pair. Darker color indicates IRESs changed, lighter color indicates IRESs not changed. (F) Effect of AS events on m6a sites in each circRNA pair. Darker color indicates m6a sites changed, lighter color indicates m6a sites not changed.

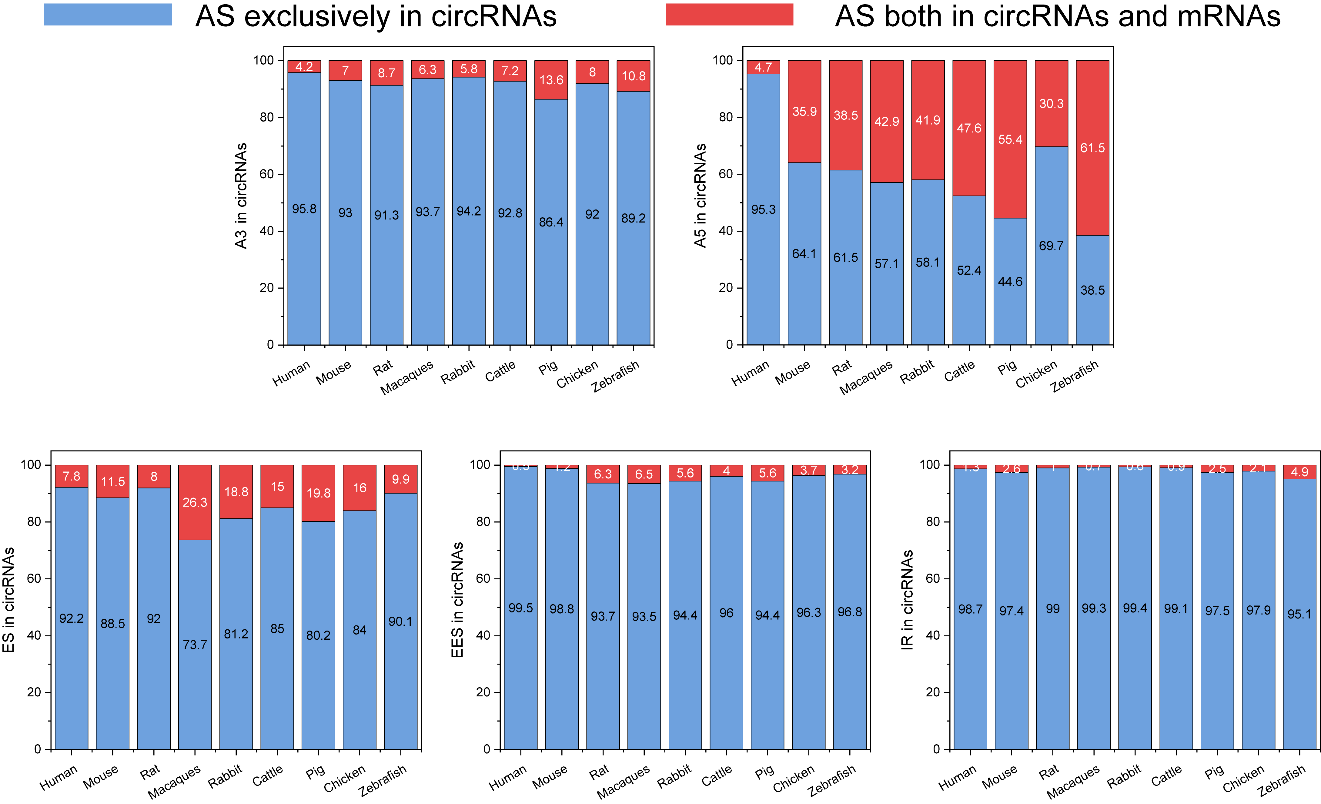

**Figure S2.** Percentage of AS events in circRNAs.

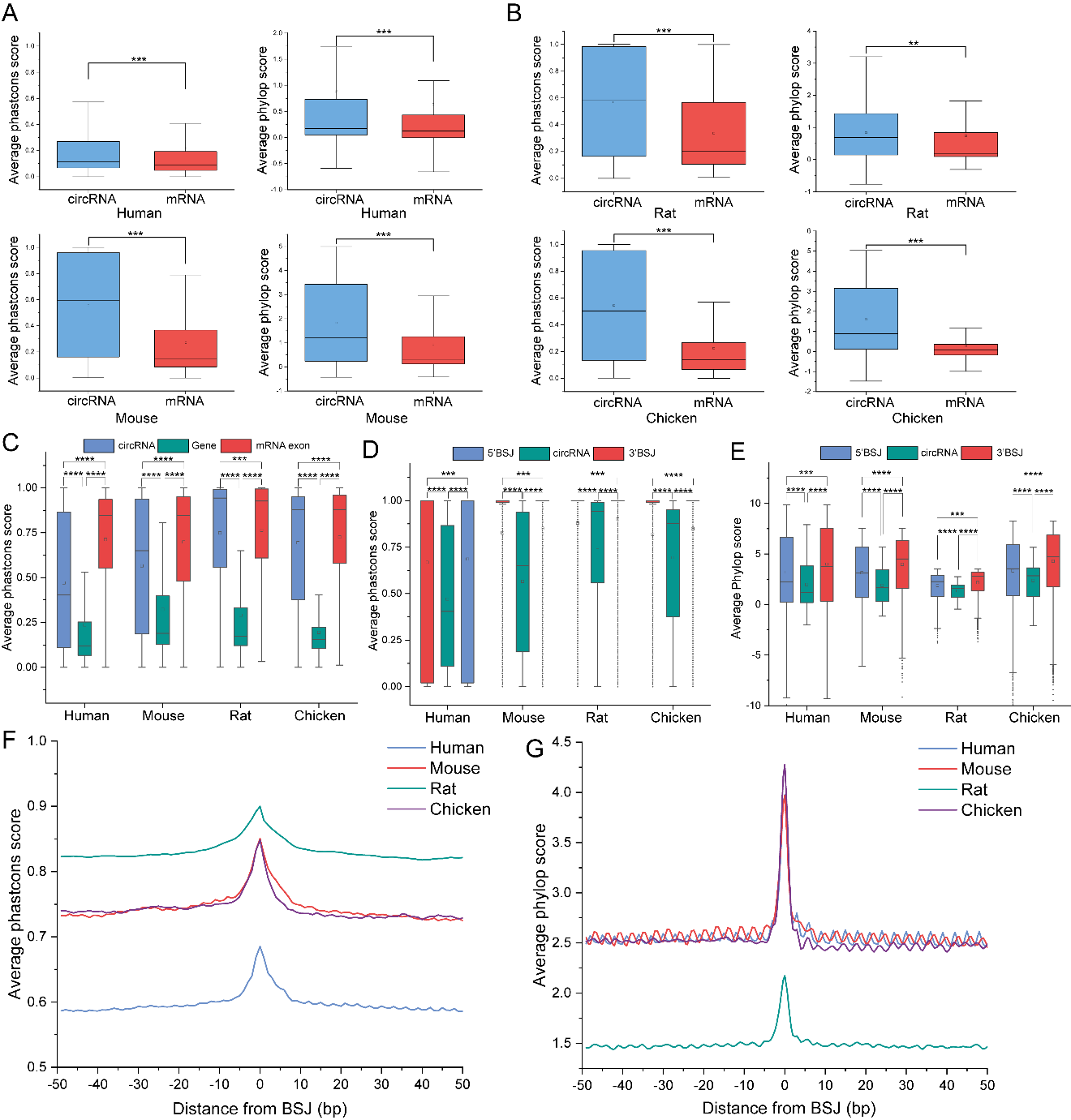

**Figure S3.** (A-B) The conservation score (Phylop and PhastCons) for circRNA-exclusive splicing fragments and mRNA-exclusive splicing fragments of four species. (Student's t-test, *:P<0.05, **:P<0.01, ***:P<0.001, ****:P<0.0001) (C) Conservation assessment (PhastCons) of circRNAs, genes and mRNA exons. (D-G) Conservation assessment (Phylop and PhastCons) near the BSJ sites.

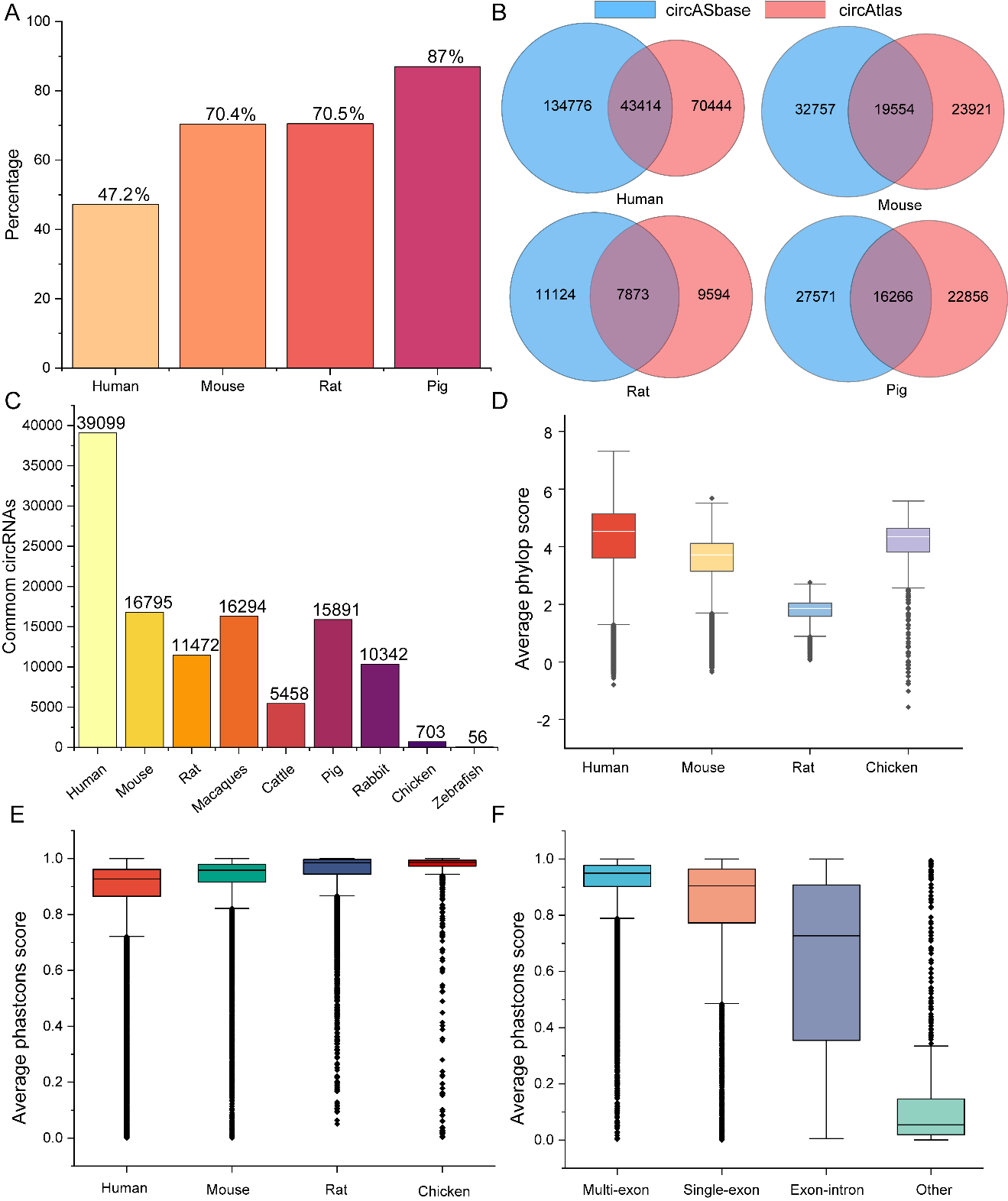

**Figure S4.** (A) Comparison of circASbase and circAtlas. Percentage of BSJs in circASbase that overlap with circAtlas. (B) Comparison of full-length circRNAs from the same BSJ. A sizable number of full-length circRNAs (Similarity is 100%) can be found in both databases. (C) Statistics of circRNAs common in two or more species. (D-F) Conservation assessment of circRNAs common in two or more species.

**Methods：**

**Assembling workflow**

1. Build BWA index

**bwa** index genome.fa

2. Aligning to the reference genome

**bwa** mem -t 12 BWAIndex xxx_1.fastq xxx_2.fastq -o xxx_results.sam

3. Identify circRNAs

**perl CIRI2.pl** -I xxx_results.sam -O xxx_circList.txt -A genes.gtf -F genome.fa

4. Build Bowtie2 index

**bowtie2-build** genome.fa genome

5. Assemble circRNA transcripts

**tophat** -o results -p 12 -G genes.gtf Bowtie2Index xxx_1.fastq xxx_2.fastq

**samtools** sort -n -o accepted_hits.sorted.bam accepted_hits.bam

**samtools** view accepted_hits.sorted.bam > accepted_hits.sorted.sam

**python CircAST.py** -L 150 -T 5 -G genes.gtf -F accepted_hits.sorted.sam -J xxx_circList.txt

**python CircFullSeq.py** -F chromfafile_folder -I CircAST_result.txt -O circRNA.fa

**SeqCIRI.pl** -i CIRI2_result -f genome.gtf -g genome.fa -out circRNA.fa

**Remove redundant sequences**

**cd-hit-est** -i input.fa -o output.fa -c 0.95 -aL 0.9

**Identify AS events**

**AS_finder.pl** -g species.gtf -c circRNA_info.result -out AS_events

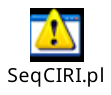

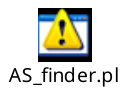
